# OrthoGuide: A database for rooting inference of orthologous genes

**DOI:** 10.1101/2025.10.20.683520

**Authors:** João V. F. Cavalcante, Gleison M. de Azevedo, Diego Marques-Coelho, Danilo O. Imparato, Mauro A. A. Castro, Rodrigo J. S. Dalmolin

## Abstract

Orthology has proven to be a valuable proxy for the study of the evolution of biological systems, such as metabolic pathways and gene regulatory networks. Genes within the same orthologous group typically share the same evolutionary history, reflecting their common ancestry. To leverage this property, several tools and databases, most notably the COG database, have been developed to support evolutionary analyses based on orthology information. Building on these resources, we previously developed the Bridge algorithm to infer the evolutionary root of genes by analyzing the distribution of orthologous groups in a phylogenetic tree. Here, we introduce OrthoGuide, a database and web application that provides rooting information for all COGs across eight model species, as inferred by the Bridge algorithm. The web application and the database are hosted at https://dalmolingroup.imd.ufrn.br/orthoguide/.

**Significance Statement:** Understanding the evolutionary origin of genes is fundamental for systems biology but is often obstructed by computationally complex bioinformatic workflows. This creates a bottleneck for many researchers. OrthoGuide directly addresses this challenge by providing a public, pre-computed database of evolutionary rooting data for all genes across eight eukaryotic model organisms. Our web application eliminates the need for user-side computation. OrthoGuide delivers instant results as well as interactive visualizations. This resource democratizes access to evolutionary analyses based on orthology information, enabling a broader scientific community to rapidly translate simple gene lists into insights about the assembly of biological systems.

## Introduction

Orthologous groups (OGs), which comprise genes descended from a single common ancestor, serve as a proxy for investigating the evolutionary history of complex biological systems. By tracing the distribution of these groups across a species tree, researchers can infer the evolutionary root, or point of origin, for each component of a system, such as a metabolic pathway or a particular trait. This approach allows for a systems-level reconstruction, moving beyond the analysis of single genes to understand how entire functional networks were assembled over time.

This methodology has yielded a variety of tools [1][2] and significant insights into the evolution of critical cellular functions. In this context, orthologous gene information has been used to unravel the evolution of neurotransmitter and apoptosis systems [3][4], to construct coevolutionary gene networks in yeast species [5] and to identify key drivers of multicellularity in animal evolution [6].

Central to this approach is the Bridge algorithm [2]. The algorithm infers a gene’s evolutionary root by systematically evaluating its phyletic pattern—the presence or absence of its orthologs—across a given species tree. A key advantage of this method is that it does not require a predefined outgroup, making it particularly well-suited for deep evolutionary analyses where a suitable outgroup may be distant or ambiguous. The Bridge algorithm operates by iteratively testing each ancestral node in the phylogenetic tree as a potential point of origin, calculating a consistency score that measures how well that root explains the observed distribution of the orthologous group through vertical descent and subsequent gene loss. The node yielding the highest score is thus inferred as the most parsimonious evolutionary root. The Bridge algorithm has been made available through an R package, GeneBridge (https://github.com/sysbiolab/GeneBridge).

Despite the power of algorithms like Bridge, their application has largely been confined to researchers with bioinformatic expertise, often requiring the use of command-line tools and package installations. This creates a technical barrier that limits the broader scientific community from leveraging these insights. OrthoGuide addresses this gap by providing a comprehensive, pre-computed database of rooting inferences generated by the Bridge algorithm, along with an intuitive accompanying web application to access it.

## Methods

### Database Implementation

The core of the OrthoGuide database consists of pre-computed evolutionary rooting information for all annotated genes of eight eukaryotic model species (Table 1). This was carried out systematically applying the Bridge algorithm, implemented in the GeneBridge R package [2], to each gene’s corresponding orthologous group.

**Table 1.**
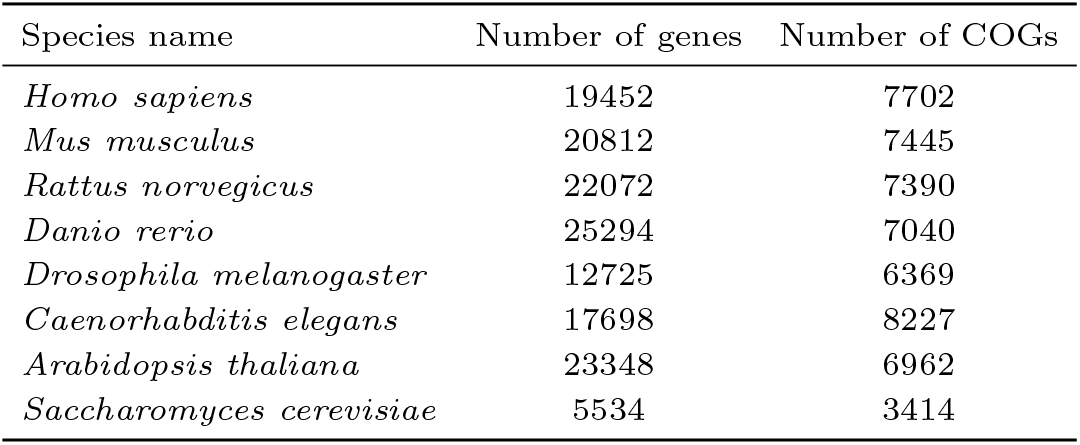
Species included in the OrthoGuide database v2.11.0.

| Species name | Number of genes | Number of COGs |
| --- | --- | --- |
| *Homo sapiens* | 19452 | 7702 |
| *Mus musculus* | 20812 | 7445 |
| *Rattus norvegicus* | 22072 | 7390 |
| *Danio rerio* | 25294 | 7040 |
| *Drosophila melanogaster* | 12725 | 6369 |
| *Caenorhabditis elegans* | 17698 | 8227 |
| *Arabidopsis thaliana* | 23348 | 6962 |
| *Saccharomyces cerevisiae* | 5534 | 3414 |

The analytical pipeline mapped all genes available in the STRING database v11.0 [7] from each of the eight model organisms to their respective Clusters of Orthologous Groups (COG) identifiers [8]. The phyletic patterns derived from these COGs served as the primary input for the rooting inference. The analysis was performed against a eukaryotic phylogenetic tree compiled using topologies from TimeTree[9] and augmented with species from the NCBI Taxonomy, containing all eukaryotes in version 11.0 of the STRING database. The tree was constructed according to a previously described methodology [3]. Each of the eight model organisms were designated as the reference species for the algorithm.

Parameters for the GeneBridge execution were set to a probability threshold of 0.5 to guide the root search and 1000 permutations. To ensure reproducibility and facilitate future updates, the entire database generation workflow was encapsulated into a containerized Nextflow pipeline [10], which allows for the seamless inclusion of new species in subsequent versions.

All organisms are organized into separate tables in the SQLite3 database, each table being named after the organism’s NCBI taxonomy identifier. All tables contain the same columns, describing the orthologous root inference (Table 2).

**Table 2.** OrthoGuide database schema, containing the columns present in each organism’s table and their data types.

| Column Name | Data Type | Description | Example |
| --- | --- | --- | --- |
| cog_id | TEXT | The Cluster of Orthologous Groups (COG) identifier. | NOG106405 |
| root | INTEGER | The numerical ID representing the last common ancestor (LCA) or root clade. | 1 |
| Dscore | REAL | The consistency score from the Bridge algorithm, measuring conformity to a vertical inheritance model. | 0.9956 |
| Statistic | REAL | The statistic value associated with the Dscore. | 8.188 |
| Pvalue | REAL | The P-value for the Dscore, indicating statistical significance. | 1.33e-16 |
| AdjPvalue | REAL | The adjusted P-value for multiple testing correction. | 8.81e-13 |
| clade_name | TEXT | The descriptive name of the root clade. | Homo |
| protein_id | TEXT | The STRING protein identifier for the specific protein. | ENSP00000490314 |
| ssp_id | INTEGER | The NCBI Taxonomy ID for the species of the protein. | 9606 |
| preferred_name | TEXT | The common gene name (e.g., HGNC symbol) used for querying. | BRCA1 |

### Web Application

The complete set of rooting results is stored in a SQLite3 database, which is accessed via a static, single-page application built with the Vue3 JavaScript framework. The use of SQLite WebAssembly enables all database queries to be executed entirely on the client-side.

OrthoGuide also provides integrated visualizations. The first is a cumulative bar graph that displays the number of genes rooted at each ancestral clade, offering an overview of the evolutionary timeline of the queried gene set. The second is an interactive protein-protein interaction (PPI) network, with interaction data retrieved from the STRING database. This feature allows users to dynamically filter the network to display only the subset of proteins that had emerged by a specific ancestral node.

## Results and Discussion

OrthoGuide is architected as a static web application that queries a SQLite database directly in the user’s browser via WebAssembly. This approach was chosen to significantly enhance performance, portability, and scalability. By executing all data operations on the client-side, the application delivers query results after an initial database load. This self-contained model also empowers other developers and research groups to easily extend or self-host their own instances of OrthoGuide.

To ensure the database itself is reproducible and extensible, it is constructed via a containerized Nextflow pipeline. This workflow automation tool guarantees that the entire data generation process is standardized, version-controlled and modifiable. Additionally, OrthoGuide’s interactive visualizations, while simple, are fully exportable as publication-ready Scalable Vector Graphics (SVG) files, allowing for lossless resizing and easy integration into academic manuscripts.

To evaluate OrthoGuide assessing the evolutionary history of a specific biological system, we analyzed the human Taste Transduction pathway from the KEGG database (hsa:map04742) [11]. This pathway, comprising 86 genes in humans, represents an excellent test case because it is a functional mosaic, containing both ancient, broadly conserved genes and more recently evolved, lineage-specific receptor families. The full gene set was submitted to the OrthoGuide application (v2.11.0) with Homo sapiens selected as the reference organism. The complete table of results can be acquired from Supplementary Table 1.

The analysis revealed a stepwise assembly of the taste transduction pathway, highlighted by two notable emergence points for major gene cohorts (Figure 1). The first and oldest major expansion event roots 18 genes to the last common ancestor (LCA) of Choanoflagellata and humans. This ancient origin is biologically meaningful, as choanoflagellates are the closest known unicellular relatives of animals. These genes, including core components of G-protein coupled receptor (GPCR) signaling and Transient Receptor Potential (TRP) channels, likely formed a molecular repertoire for environmental chemosensation, which is consistent to what is known about this clade [12].

**Fig. 1.**
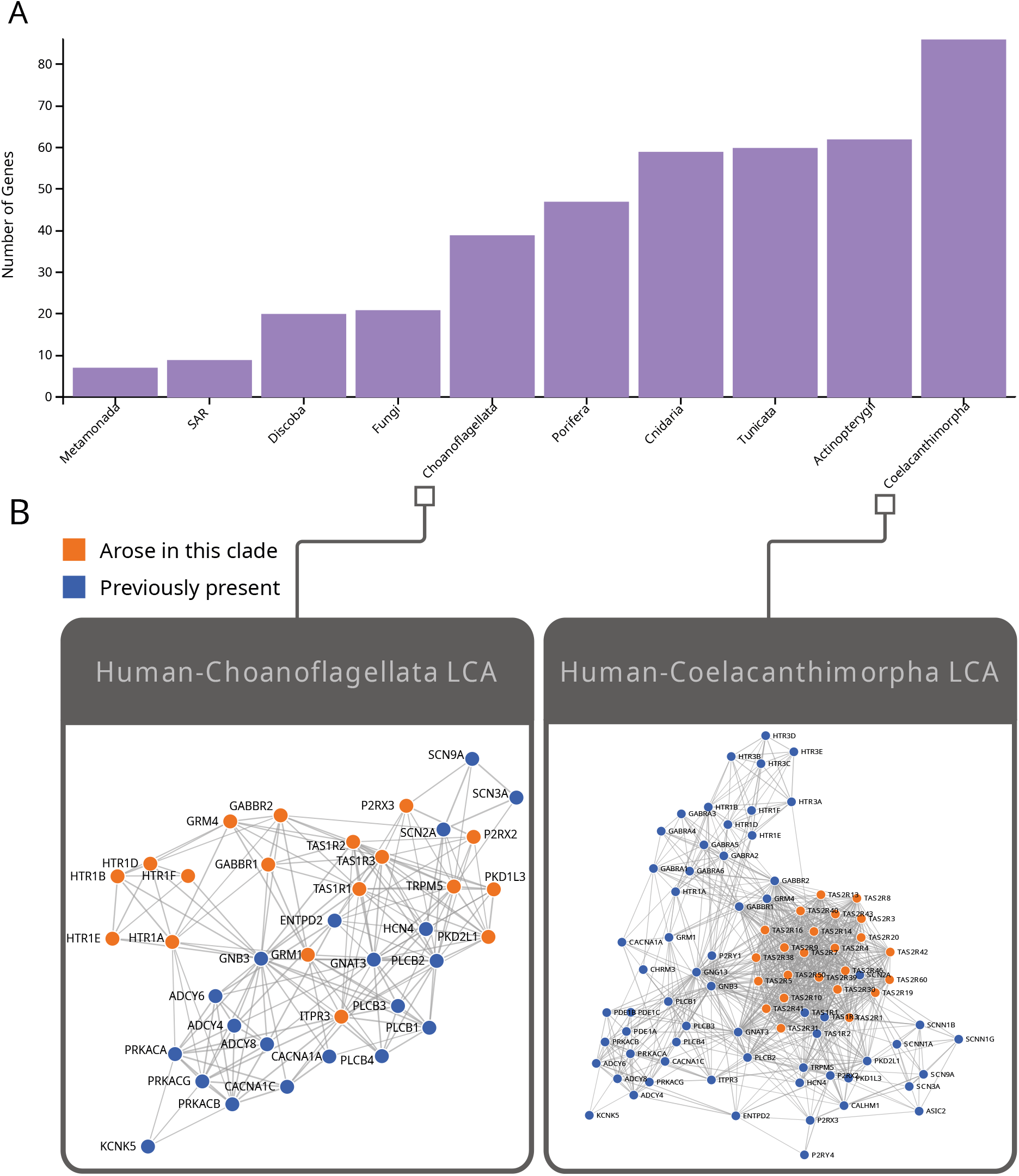
Visualizations from OrthoGuide for genes from the Taste transduction pathway. **A**: The cumulative number of rooted genes at each ancestral point in the species tree for this set. **B**: Protein-protein interaction networks considering the cumulative presence of rooted nodes from each root clade. Nodes rooted at each point are shown in orange while nodes rooted in earlier clades are shown in blue.

A second expansion event occurred later, with 24 genes emerging in the LCA of Coelacanthimorpha and humans. This diversification is dominated by the evolution of bitter taste perception, mediated by the Taste Receptor Type 2 (T2R) family. The coelacanth is a critical node in vertebrate evolution and is noted for being the first fish lineage to possess an extensive repertoire of bitter taste receptors, likely as an adaptation to detect toxins in a more complex diet [13] [14]. This evolutionary finding is also clearly visualized in our results as the emergence of a highly connected module of proteins belonging to the T2R family.

In summary, this evaluation demonstrates OrthoGuide’s ability to uncover a broader evolutionary narrative of a biological system. The results not only pinpoint key evolutionary events but also place them in their proper biological context.

## Conclusions

OrthoGuide provides a systems biology-based platform for the evolutionary analysis of genes and biological pathways, designed to identify the evolutionary origins of functionally associated genes. The core advantage of OrthoGuide over existing analytical workflows is its accessibility; it provides pre-computed rooting information generated by the Bridge algorithm, which infers the evolutionary age of a gene based on the phyletic distribution of its orthologous group. OrthoGuide eliminates technical obstacles by presenting the results in an intuitive, query-based web application, enhanced by interactive visualizations that allow users to explore the stepwise assembly of a system’s components over time. The OrthoGuide database and web application thus serve as a well-documented and easy-to-use resource, developed to help bioinformaticians and biologists convert their gene lists into biological insights about the evolutionary history of their systems of interest.

It is important, however, to acknowledge the limitations inherent to this pre-computed approach. The accuracy of OrthoGuide’s inferences is dependent on the quality and comprehensiveness of the curated orthology database it is built upon. Any inaccuracies in the underlying orthology assignments will naturally propagate into our results. Furthermore, because the database is static, users cannot infer new evolutionary scenarios using custom phylogenetic trees or for species not yet included in our dataset. This trade-off is by design, as it enables the platform’s primary strength: delivering instantaneous, resource-free analysis for the end-user. Therefore, OrthoGuide is positioned not as a replacement for dynamic analysis pipelines, but as a tool for rapid hypothesis generation and initial data exploration within its defined evolutionary context.

## Software and Data Availability

GeneBridge is released under the Artistic-2.0 license. The source code for GeneBridge is available at https://github.com/sysbiolab/GeneBridge. OrthoGuide is released under the Apache 2.0 license and its source code and database are available at https://github.com/dalmolingroup/orthoguide/. The OrthoGuide application can be found at https://dalmolingroup.imd.ufrn.br/orthoguide/.

## Competing interests

No competing interest is declared.

## Author contributions statement

J.V.F.C, G.M.A. and D.M.C conceived the database and application, J.V.F.C. developed the database and application, D. O. I. assisted in the database development, J.V.F.C. and G.M.A. conducted the application analysis, J.V.F.C, M.A.A.C and R.J.S.D wrote the manuscript. All authors reviewed and approved the final manuscript.

## Acknowledgments

J.V.F.C would like to thank Patrick Terrematte for valuable feedback during the application development. This study was financed by the Brazilian Agency Coordination for the Improvement of Higher Education Personnel (CAPES – Portuguese: Coordenação de Aperfeiçoamento de Pessoal de Nível Superior - Brasil (CAPES)-(project numbers 88881.692896/2022-01 and 88887.002655/2024-00), National Council of Technological and Scientific Development (CNPq – Portuguese: Conselho Nacional de Desenvolvimento Científico e Tecnológico (CNPq) (project number 312305/2021-4 and 305019/2025-2), PROPESQ-UFRN. We would also like to thank NPAD / UFRN for computational resources.

## Notes

https://dalmolingroup.imd.ufrn.br/orthoguide/

### Competing Interest Statement

The authors have declared no competing interest.

